# Age-specific activation of microglial genes in a mouse model of AD

**DOI:** 10.1101/2022.10.28.514172

**Authors:** Haoyang Zhou, Yizhou Yu

## Abstract

Alzheimer’s disease (AD) is a neurodegenerative disease that increases its prevalence with age. AD is characterized by a progressive loss of cognitive function, leading to dementia. The conventional pathological hallmarks of AD include deposition of amyloid plaque, aggregation of neurofibrillary tangles, and sustained inflammation. Recent studies have confirmed that immune responses play an important role in AD progression. Single-cell RNA-sequencing (RNA-seq) studies of the AD mouse model identified the disease-associated microglia (DAM) population during AD progression, where the pro-inflammatory activity was mediated by Trem2. However, the time points and nature of DAM activation remain largely unknown. Here, we compared the time-result bulk sequencing andsingle nuclei RNA sequencing data in a mouse model of AD. We identified an age-dependent activation of Trem2 targeted immune-related genes. We showed that Trem2 affected DAM activation at an early stage (4-8 months) rather than the late stage (12-16 months). Our results highlight an age-dependent change in the Trem2-linked immune response in AD.

## INTRODUCTION

Alzheimer’s disease (AD) is an irreversible neurodegenerative disease. The prevalence of AD increases with age, with around 0.6% of the population aged between 65 and 69, even 8.4% after 85 years old [1–4]. The primary symptom of AD is cognitive dysfunction which eventually leads to dementia [5]. Disease-modifying treatments of AD are currently limited [6].

In order to find disease-modifying therapeutics for AD, it is thus a necessity to explore the molecular pathways involved in AD. Neuroinflammation has been extensively studied as a hallmark of AD. Microglia has been shown to remove amyloid-β and damaged brain cells via phagocytosis. However, chronic stimulation of microglia by Aβ or other misfolded proteins may lead to prolonged inflammation [2,14]. Post-mortem studies revealed that young patients with systemic inflammation have similar morphological changes of microglia to those in dementia patients [15]. In addition, asymptomatic patients show no relevance between such microglial morphological changes and Aβ load, indicating neuroinflammation may be an independent or even prerequisite pathological event of AD amyloid plaque [15,16]. Genome-wide analyses have shown that the most differentially expressed genes between AD and control are immune genes rather than neural genes [17]. Furthermore, single-cell transcriptome studies have revealed that these AD-related immune response genes, also termed disease-associated microglia (DAM) genes, can be activated by a key immunological gene, Trem2 (Triggering Receptor Expressed on Myeloid Cells 2), expressed in disease-specific microglial populations [17,18]. Several mutations of Trem2, such as R47H and R62H, have been identified to increase late-onset AD risk by 2 to 4 folds [19,20]. Consistently, Trem2 KO exacerbates the AD phenotypes in 5xFAD mice, increasing Aβ load and neuron loss. However, Trem2 KO has also been shown to attenuate the expression of inflammatory cytokine transcripts in the same 5xFAD model [21]. The role of DAM in AD and whether the gene expression profile change in DAM with respect to age remains intriguing. A major roadblock in identifying prevalent immune-related drug targets is the detailed understanding of when and where DAM and neuroinflammation are engaged in AD.

Here, we focused on age-related gene expression in mouse model of AD (5xFAD). We combined bulk RNA-seq and single-cell RNA-seq to analyze age-related changes and microglial activation. We found that inflammatory gene activation is age-specific. Through analysis of single-nuclei RNA-seq data, we found a disease-specific population in early rather than late-stage. Our results suggest that AD treatment targeting immune responses should be tuned to disease stages.

## METHODS

### Data collection

GSE168137 [24] was used for bulk RNA-seq data analysis to understand the specific genes and associated pathways of AD. Samples were collected from the hippocampus and the cortex tissue in 5xFAD mice and wildtypes at 4, 8, 12, and 18 months old. Data were aligned to mouse references mm10 and gene expressions were normalized to TPM (Transcripts per Million mapped reads). GSE140510 and GSE140399 [25] were analyzed for single-cell RNA-seq to show details in microglia and other glial populations. GSE140510 was from cortex in mice at 7 months old (considered as early stage in AD) and GSE140399 was from mice at 15 months old (late stage), with each containing 4 genotypes, including wildtype, 5xFAD, Trem2 KO, and 5xFAD; Trem2 KO. The UMI count matrix was collected for downstream analysis [25].

### Data preprocessing

In bulk RNA-seq data, lowly expressed genes were filtered out and gene expression matrices were finalized by creating a threshold of TPM≥1 and expressed in at least 5 replicates. Further, the data were quantile normalized using the R package, Limma [26]. In single-cell RNA-seq data, samples with high mitochondria percentage and low number of expressed genes were filtered out. This analysis was conducted by R package, Seurat [27].

### Principal component analysis (PCA)

PCA is used for dimension reduction to isolate the most important principal components that account for the highest variance. PCA is conducted here to visualize batch effects and technical effects in RNA-seq experiments. Bulk gene expression matrix was first log2 transformed and fed into R function prcomp, with center equals to true and scale equals to false.

### Differential gene expression analysis

In order to understand the differentially expressed genes in AD, read counts of genes are compared between disease samples and wild type. A gene is declared differentially expressed when the difference is significant (FDR<0.05). ExactTest, from EdgeR package [28], was used to conduct differential gene expression analysis.

### Time course differential analysis

Time course differential analysis across brain regions, time points or genotypes was done by masigPro [29]. Rsq=0.7 and T.fit α=0.05 were defined to cluster differentially expressed genes and isolate AD-related genes, brain region-specific genes, and aging-related genes.

### Gene ontology analysis

Gene ontology is an effort to unify the terminology of gene function. To understand the specific pathway of the upregulated and downregulated genes, gene lists were fed into Metascape [30] to conduct gene ontology analysis with input species as mouse. Enriched terms were selected based on hypergeometric tests (BH-corrected p-value<0.05).

### Single-cell clustering

Single-cell clustering was conducted by R package, Seurat [27]. The dimension reduction was conducted with the UMAP algorithm to observe the cell type clustering based on their gene markers. Mixed population and doublets were removed and cell type-specific markers and differential markers between treatments/genotypes were found.

### Normalization for relative enrichment

Relative enrichment of each cell type within a cluster is calculated with the following equation.

## RESULTS

### Upregulation of immune response genes in AD

In order to understand the temporal pattern of neuroinflammation in AD, we analysed gene expression data in different brain regions and ages of a mouse model of AD. We collected published bulk RNA-seq samples from the cortex and hippocampus in both 5xFAD and WT mice at the age of 4, 8, 12, and 18 months respectively [25]. Using principal component analysis, we found that the gene expression signatures from the hippocampus and the cortex were separated by time or age (Figure 1A and B). In order to isolate genes linked to AD, we analysed the spararate clusters. K-means clustering was used to further classify differentially expressed genes, resulting in 8 clusters based on their expression profiles. We observed gene expression patterns that correlate with age, brain region or genotype (Figure 1C). We found a cluster of genes specific to the effect of the genetic mutation introduced to model AD (FAD). We performed an ontology analysis on the genes in the FAD cluster and found that a signature of innate immunity, cell activation and inflammatory response. To confirm the activation of the immune response, we compared the differentially expressed genes in our dataset to a list of DAM genes [31] to observe how many immune-related genes were activated. In both hippocampus and cortex, there were more genes upregulated in the 5xFAD model compared to the WT (Supplementary Table 1, available in our GitHub repository). We found that DAM genes are highly upregulated in the early stage of AD progression, while their proportions decrease as age increases. While such gene expression patterns remain consistent in the cortex and the hippocampus, the effect on the hippocampus can be observed at a relatively earlier time point.

**Figure 1:**
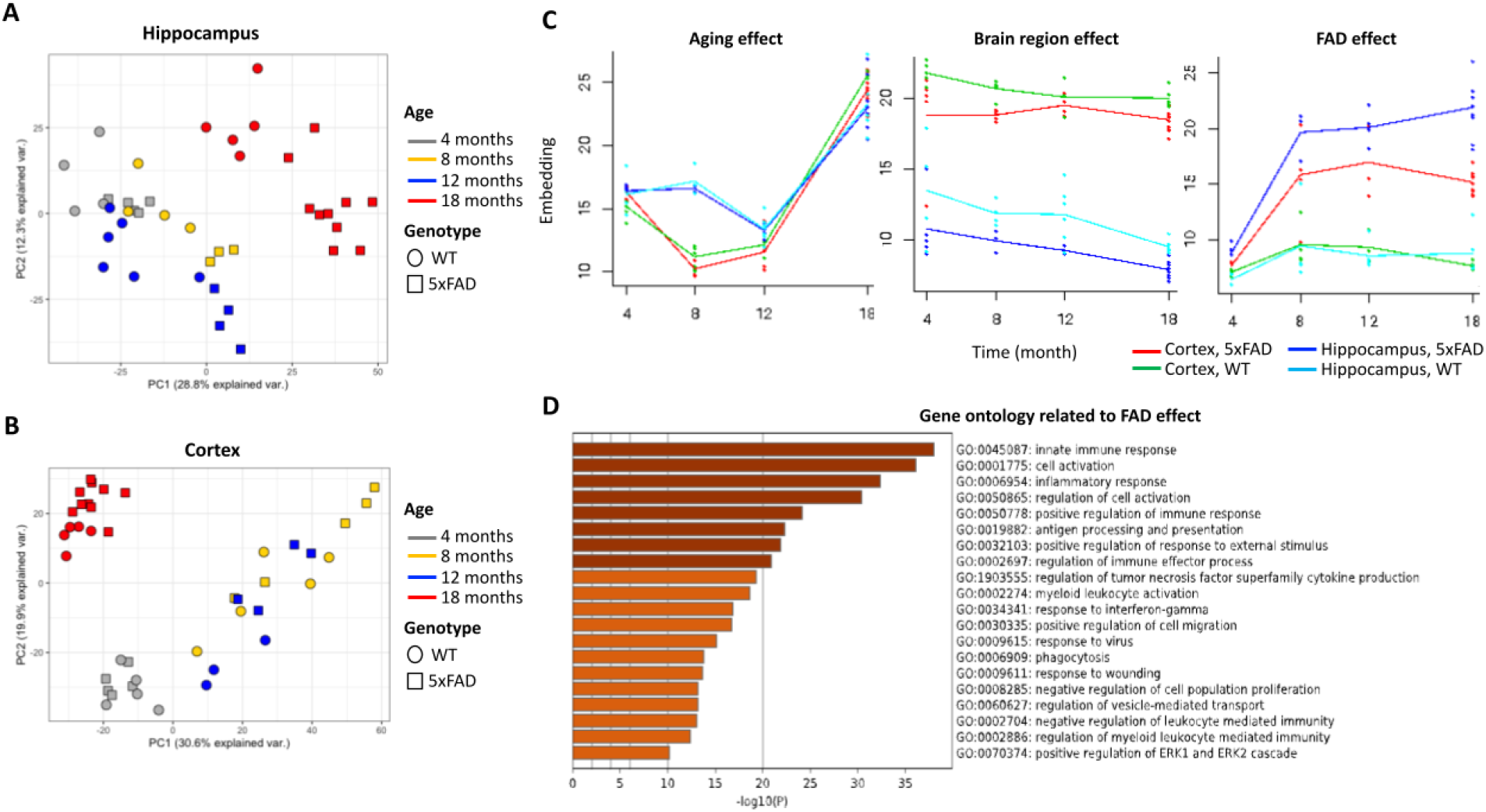
Upregulated genes and relevant GO terms in FAD vs WT mice. (A & B) Principal components of RNA-seq data from 5xFAD and WT mice in the hippocampus (A) and cortex (B) across 4 time points. The time point was represented by different colors and shapes denoting the genotype of mice. C. Gene expression clusters linked to age, brain region and genotype (FAD). E. Ontology associated with the cluster related to FAD. Significant terms were highlighted in darker brown compared to light brown, and ranked by significance level.

### Trem2-dependent enrichment of microglia in an early stage during disease progression

Our age-related analysis of AD in a mouse model showed a link in immune activation. Since multiple types of cells can mediate neuroinflammation, we analysed single cell sequencing data in young and old 5xFAD and control mice. We compared the transcriptome in different cell types between 7 months and 15 months, with/without Trem2 [25,32]. The UMAP algorithm was applied to reduce dimensions on the single nuclei data, and cell clusters were generated using the data at 7 months and 15 months (Figure 2A and B). By comparing similar numbers of recovered nuclei, we found more neuron diversity (more neuronal clusters) in 15 months than 7 months. Further, we observed lower proportions of glial cells at 15 months compared to the 7 months data. In order to analyse the changes in cell types, we performed enrichment analysis. We found that the microglia were more specifically enriched in FAD genotype at 7 months, but not in other genotypes (WT, WT Trem2 KO, and FAD Trem2 KO) (Figure 2C). At 15 months, this enrichment was not observed (Figure 2D). In 15 months, both Trem2 KO and FAD Trem2 KO have a lower enrichment in microglia, while aged WT mice and FAD mice have similar levels of enrichment. Our results suggest that microglia activationcould be dependent on Trem2. While Trem2 KO 5x FAD mice have accelerated AD phenotypes [2], our results suggest that microglia activation may not be always associated with aggravation of AD. We therefore sought to investigate DAM-specific gene expression profile.

**Figure 2:**
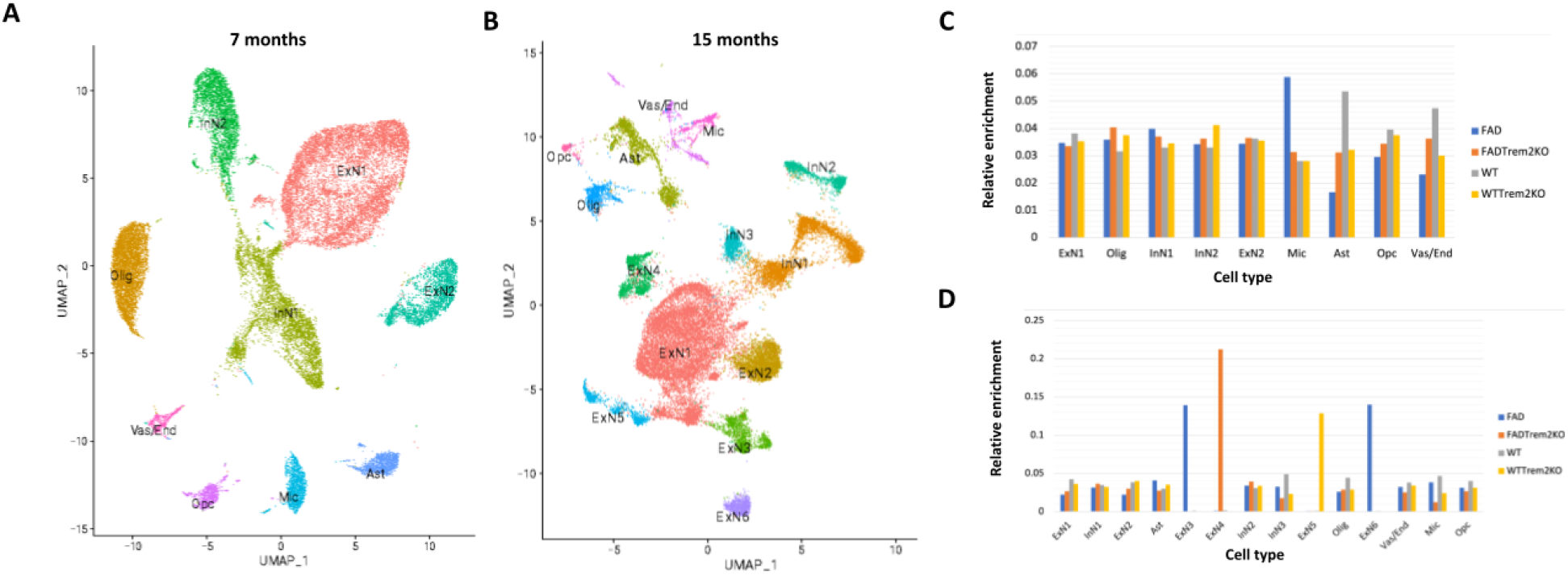
Trem2 and age-dependent increase in microglial cells in a mouse model of AD. A&B. Cell type clustering based upon single-nuclei RNA-seq data at the early stage, 7 months (A) and late stage, 15 months (B). Nuclei were labeled by cell type and genotypes in UMAP, number and proportion of nuclei for each cell type were listed. The abbreviations are: ExN - excitatory neuron, InN - inhibitory neuron, Olig – oligodendrocyte, Ast – astrocyte, Mic – microglia, Opc-Oligodendrocyte progenitor cell, Vas/End – vascular and endothelial cells. C&D. Trem2-dependent enrichment of microglial cells in young (7 months) 5xFAD mice. The number of nuclei was normalized by the number of cells in the cell type and the number of cells in each genotype to obtain relative enrichment.

To observe how specific DAM genes changed in response to Trem2 KO in early and late stages, first looked at cell clusters in young (Figure 3A) and older mice (Figure 3B). We observed distinct clusters in young mice. Additionally, Trem2KO shifted the FAD cluster towards controls, further supporting our observations of a Trem2-dependent effect in young mice models of AD.

**Figure 3:**
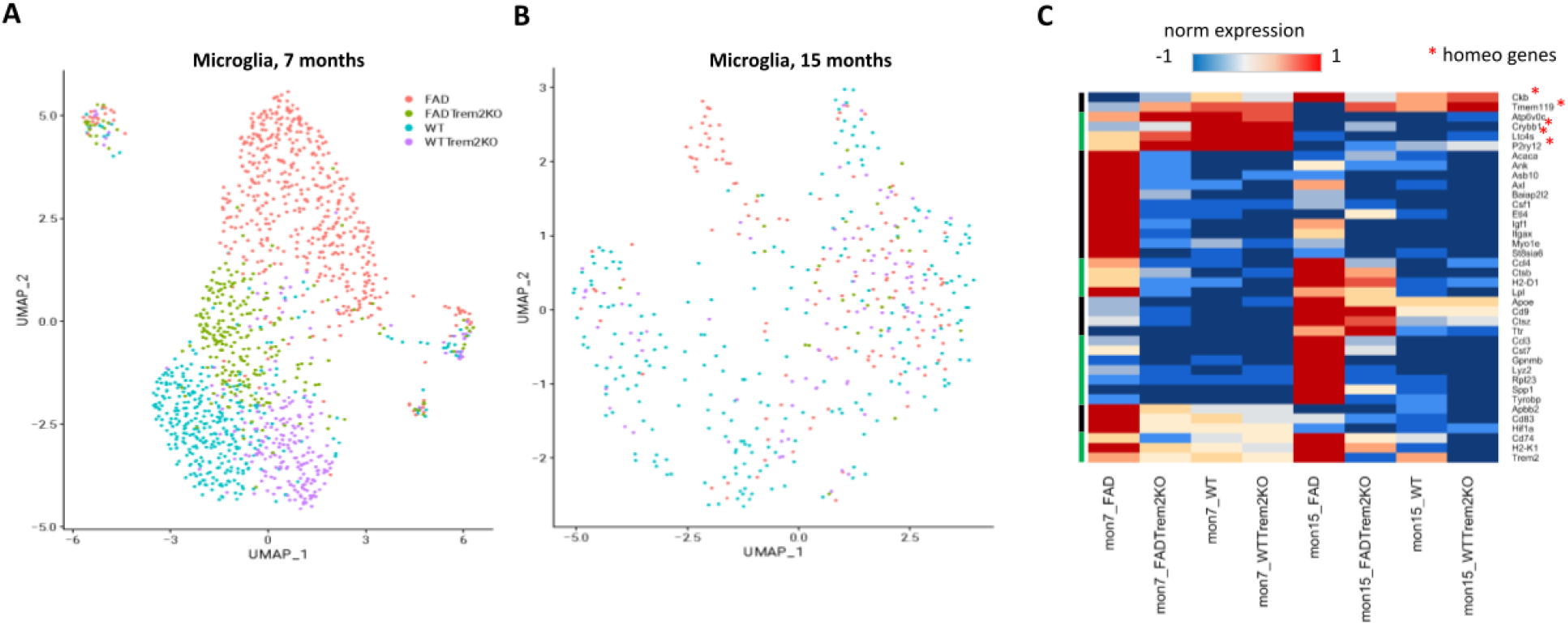
Age-specific comparison of microglial cells in models of AD. (A & B) Sub Clustering of microglia nuclei in 7 months (A) and 15 months (B). Nuclei were colored by genotypes. C. Heatmap of gene expression of 38 genes that were differentially expressed between FAD and WT and known as DAM or homeostatic microglia markers. K-means clustering was performed to group them based on expression patterns across genotypes and time points, labeled on the left side of the heatmap. Gene expression TPM was the first quantile normalized and then transformed between −1 and 1 (blue, low expression; red, high expression). Homeostatic microglia markers were labeled with red asterisk.

We did not observe any clear evidence of disease-specific clustering of microglia the microglia of older mice (Figure 3B), suggesting that the effect of FAD on microglia is temporally restricted to younger mice.

Microglia contribute to the maintenance of brain homeostasis, but lose homeostatic function during neurodegenerative disorders. We then sought to gain insight into the gene expression of these microglia, and their link to DAM and their function in maintaining brain homeostasis. We extracted differentially expressed genes between FAD and WT (Figure 5A) in both 7 months and 15 months, and then compared these genes with published DAM marker list and homeostatic microglia marker list [31]. Then we performed k-means clustering on these intersected genes and obtained 8 clusters, showing different gene expression profiles in early and late stages among various genotypes (Figure 3C). Of the 38 common genes, only 5 were homeostatic microglia markers while the rest were DAM genes. Homeostatic microglia markers were only upregulated only at 7 months, not 15 months. This suggest that the upregulation of microglia in young mice could confer a protective effect.

On the other hand, most DAM genes seemed to be highly Trem2 dependent in both stages, suggesting the potentially critical role for Trem2 in DAM activation. More specifically, DAM genes could be separated into two distinct categories based on age (Figure 5C). This finding suggested microglia could have different roles in the early and late stages of the disease. Linking this analysis back to our clustering of microglia nuclei, we found a disease-specific population in the early stage rather than the late stage of disease progression. Taken together, our results suggest different functions of microglia at the different stages of the AD-like pathology modelled in mice. This difference could be due to a distinct microglial cluster.

## DISCUSSION

Consistent with previous studies, we showed that more DAM genes become activated with AD progression. Using bulk and single-cell analysis, we explored the characteristics of DAM further. Intriguingly, we observed that DAM showed distinct profiles in the early and late stages of AD. This indicated the timepoint-dependent mediation of DAM genes in disease-associated activation. We suggest that DAM initially plays a protective role but switches to a chronic inflammatory role later which damages neurons in AD. Furthermore, by classifying genes associated with the early and late stages, we found that more genes in the late stage are inflammation related. Such results supported the similar model proposed by Onuska [33], in which DAM progressively becomes more inflammation related. Finally, we observed that there were clearer boundaries of microglia between different genotypes in the early stage than in the late stages, suggesting that early-stage DAM had more distinct characteristics compared to WT counterparts.

In previous studies, Trem2 was identified to be a key player in DAM activation [17,18]. Here, we confirmed that Trem2 was highly important in DAM activation. Unexpectedly, 5xFAD Trem2 KO samples were characteristically closer to WT than their disease counterparts. However, Trem2 KO in 5xFAD mouse model is known to accelerate the AD phenotype compared with 5xFAD. One potential explanation is Trem2 is a trigger to DAM, yet the downstream effect of DAM is dependent on other regulatory factors and not Trem2. In contrast, Trem2 seems to play a different role in astrocytes, making the WT Trem2 KO samples closer in characteristic to the disease sample. This is similar to the finding that the loss of Trem2 in A1 reactive astrocytes is associated with cerebral amyloid angiopathy [34]. Further, our results showed that some previously identified Trem2 independent markers seemed Trem2 dependent in later stages.

This study provides insights into the development of novel AD treatment. Firstly, targeting earlier stages of AD might be more viable compared to later stages due to more distinct characteristics in microglia. This will avoid unnecessary off-target effects on normal aged microglia. Secondly, when developing treatment, interactions with other neuroglia need consideration. Those cells are also developing their own disease-specific clusters in our results, meaning they also contribute to AD. Furthermore, data from Trem2 knockout showed that other glial cells might also respond to microglia-specific treatments, necessitation a more precise study of cell-cell interactions.

In conclusion, we further characterized the DAM and found that it is time-point dependent. We confirmed that Trem2 plays a significant role in DAM activation.

## ACKNOWLEDGEMENTS

I would like to thank Dr. Jiang and Dr. Tian for their insightful suggestions on this manuscript.

## DATA AND CODE AVAILABILITY

All data and code, as well as Supplementary Table 1 will be available in our GitHub repository: https://github.com/izu0421/AD_seq_microglia

## REFERENCES

[1] Hebert, Liesi E. “Age-Specific Incidence of Alzheimer’s Disease in a Community Population.” JAMA: The Journal of the American Medical Association, vol. 273, no. 17, 1995, p. 1354., doi:10.1001/jama.1995.03520410048025.

[2] Long, Justin M., and David M. Holtzman. “Alzheimer Disease: An Update on Pathobiology and Treatment Strategies.” Cell, vol. 179, no. 2, 2019, pp. 312–339., doi:10.1016/j.cell.2019.09.001.

[3] Drew, Liam. “An Age-Old Story of Dementia.” Nature, vol. 559, no. 7715, 2018, doi:10.1038/d41586-018-05718-5.

[4] Special Report - Alzheimer’s Association. www.alz.org/media/Documents/alzheimers-facts-and-figures-special-report-2021.pdf.

[5] “What Is Alzheimer’s Disease?” Centers for Disease Control and Prevention, Centers for Disease Control and Prevention, 26 Oct. 2020, www.cdc.gov/aging/aginginfo/alzheimers.htm.

[6] Yiannopoulou, Konstantina G., and Sokratis G. Papageorgiou. “Current and Future Treatments for Alzheimer’s Disease.” Therapeutic Advances in Neurological Disorders, vol. 6, no. 1, 2012, pp. 19–33., doi:10.1177/1756285612461679.

[7] Tampi, Rajesh R, et al. “Aducanumab: Evidence from Clinical Trial Data and Controversies.” Drugs in Context, vol. 10, 2021, pp. 1–9., doi:10.7573/dic.2021-7-3.

[8] “How Is Alzheimer’s Disease Treated?” National Institute on Aging, U.S. Department of Health and Human Services, www.nia.nih.gov/health/how-alzheimers-disease-treated.

[9] Commissioner, Office of the. “FDA Grants Accelerated Approval for Alzheimer’s Drug.” U.S. Food and Drug Administration, FDA, www.fda.gov/news-events/press-announcements/fda-grants-accelerated-approval-alzheimers-drug.

[10] Knopman, David S., et al. “Failure to Demonstrate Efficacy of Aducanumab: An Analysis of the Emerge and Engage Trials as Reported by Biogen, December 2019.” Alzheimer’s & Dementia, vol. 17, no. 4, 2020, pp. 696–701., doi:10.1002/alz.12213.

[11] Congdon, Erin E., and Einar M. Sigurdsson. “Tau-Targeting Therapies for Alzheimer Disease.” Nature Reviews Neurology, vol. 14, no. 7, 2018, pp. 399–415., doi:10.1038/s41582-018-0013-z.

[12] Sabbagh, Marwan Noel, and Jeffrey Cummings. “Open Peer Commentary to ‘Failure to Demonstrate Efficacy of Aducanumab: An Analysis of the Emerge and Engage Trials as Reported by Biogen December 2019.’” Alzheimer’s & Dementia, vol. 17, no. 4, 2020, pp. 702–703., doi:10.1002/alz.12235.

[13] Sayas, Carmen Laura. “Tau-Based Therapies for Alzheimer’s Disease: Promising Novel Neuroprotective Approaches.” Neuroprotection in Autism, Schizophrenia and Alzheimer’s Disease, 2020, pp. 245–272., doi:10.1016/b978-0-12-814037-6.00005-7.

[14] Bachiller, Sara, et al. “Microglia in Neurological Diseases: A Road Map to Brain-Disease Dependent-Inflammatory Response.” Frontiers in Cellular Neuroscience, vol. 12, 2018, doi:10.3389/fncel.2018.00488.

[15] Streit, Wolfgang J., et al. “Dystrophic (Senescent) Rather than Activated Microglial Cells Are Associated with Tau Pathology and Likely Precede Neurodegeneration in Alzheimer’s Disease.” Acta Neuropathologica, vol. 118, no. 4, 2009, pp. 475–485., doi:10.1007/s00401-009-0556-6.

[16] Leng, Fangda, and Paul Edison. “Neuroinflammation and Microglial Activation in Alzheimer Disease: Where Do We Go from Here?” Nature Reviews Neurology, vol. 17, no. 3, 2020, pp. 157–172., doi:10.1038/s41582-020-00435-y.

[17] Landel, Véréna, et al. “Temporal Gene Profiling of the 5XFAD Transgenic Mouse Model Highlights the Importance of Microglial Activation in Alzheimer’s Disease.” Molecular Neurodegeneration, vol. 9, no. 1, 2014, p. 33., doi:10.1186/1750-1326-9-33.

[18] Keren-Shaul, Hadas, et al. “A Unique Microglia Type Associated with Restricting Development of Alzheimer’s Disease.” Cell, vol. 169, no. 7, 2017, doi:10.1016/j.cell.2017.05.018.

[19] Guerreiro, Rita, et al. “trem2 Variants in Alzheimer’s Disease.” New England Journal of Medicine, vol. 368, no. 2, 2013, pp. 117–127., doi:10.1056/nejmoa1211851.

[20] Gratuze, Maud, et al. “New Insights into the Role of Trem2 in Alzheimer’s Disease.” Molecular Neurodegeneration, vol. 13, no. 1, 2018, doi:10.1186/s13024-018-0298-9.

[21] Wang, Yaming, et al. “Trem2 Lipid Sensing Sustains the Microglial Response in an Alzheimer’s Disease Model.” Cell, vol. 160, no. 6, 2015, pp. 1061–1071., doi:10.1016/j.cell.2015.01.049.

[22] Liddelow, Shane A., et al. “Neurotoxic Reactive Astrocytes Are Induced by Activated Microglia.” Nature, vol. 541, no. 7638, 2017, pp. 481–487., doi:10.1038/nature21029.

[23] Dong, Yuan, et al. “Drug Development for Alzheimer’s Disease: Microglia Induced Neuroinflammation as a Target?” International Journal of Molecular Sciences, vol. 20, no. 3, 2019, p. 558., doi:10.3390/ijms20030558.

[24] Forner, Stefania, et al. “Systematic Phenotyping and Characterization of the 5xfad Mouse Model of Alzheimer’s Disease.” 2021, doi:10.1101/2021.02.17.431716.

[25] Zhou, Yingyue, et al. “Human and Mouse Single-Nucleus Transcriptomics Reveal trem2-Dependent and TREM2-Independent Cellular Responses in Alzheimer’s Disease.” Nature Medicine, vol. 26, no. 1, 2020, pp. 131–142., doi:10.1038/s41591-019-0695-9.

[26] Ritchie, Matthew E., et al. “Limma Powers Differential Expression Analyses for RNA-Sequencing and Microarray Studies.” Nucleic Acids Research, vol. 43, no. 7, 2015, doi:10.1093/nar/gkv007.

[27] Hao, Yuhan, et al. “Integrated Analysis of Multimodal Single-Cell Data.” Cell, vol. 184, no. 13, 2021, doi:10.1016/j.cell.2021.04.048.

[28] Robinson, M. D., et al. “Edger: A Bioconductor Package for Differential Expression Analysis of Digital Gene Expression Data.” Bioinformatics, vol. 26, no. 1, 2009, pp. 139–140., doi:10.1093/bioinformatics/btp616.

[29] Nueda, M. J., et al. “Next Masigpro: Updating Masigpro Bioconductor Package for RNA-Seq Time Series.” Bioinformatics, vol. 30, no. 18, 2014, pp. 2598–2602., doi:10.1093/bioinformatics/btu333.

[30] Zhou, Yingyao, et al. “Metascape Provides a Biologist-Oriented Resource for the Analysis of Systems-Level Datasets.” Nature Communications, vol. 10, no. 1, 2019, doi:10.1038/s41467-019-09234-6.

[31] Sobue, Akira, et al. “Microglial Gene Signature Reveals Loss of Homeostatic Microglia Associated with Neurodegeneration of Alzheimer’s Disease.” Acta Neuropathologica Communications, vol. 9, no. 1, 2021, doi:10.1186/s40478-020-01099-x.

[32] Ulland, Tyler K., and Marco Colonna. “Trem2 — a Key Player in Microglial Biology and Alzheimer Disease.” Nature Reviews Neurology, vol. 14, no. 11, 2018, pp. 667–675., doi:10.1038/s41582-018-0072-1.

[33] Onuska, Kate M. “The Dual Role of Microglia in the Progression of Alzheimer’s Disease.” The Journal of Neuroscience, vol. 40, no. 8, 2020, pp. 1608–1610., doi:10.1523/jneurosci.2594-19.2020.

[34] Taylor, Xavier, et al. “A1 Reactive Astrocytes and a Loss of TREM2 Are Associated with an Early Stage of Pathology in a Mouse Model of Cerebral Amyloid Angiopathy.” Journal of Neuroinflammation, vol. 17, no. 1, 2020, doi:10.1186/s12974-020-01900-7.

